# Sex-specific impact of patterns of imageable tumor growth on survival of primary glioblastoma patients

**DOI:** 10.1101/325464

**Authors:** Paula Whitmire, Cassandra R. Rickertsen, Andrea Hawkins-Daarud, Eduardo Carrasco, Julia Lorence, Gustavo De Leon, Lee Curtin, Spencer Bayless, Kamala Clark-Swanson, Noah C. Peeri, Christina Corpuz, Christine Paula Lewis-de los Angeles, Bernard R. Bendok, Luis Gonzalez-Cuyar, Sujay Vora, Maciej Mrugala, Leland S. Hu, Lei Wang, Alyx Porter, Priya Kumthekar, Sandra K. Johnston, Kathleen M. Egan, Robert Gatenby, Peter Canoll, Joshua B. Rubin, Kristin R. Swanson

## Abstract

**Background:** Sex is recognized as a significant determinant of outcome among glioblastoma patients, but the relative prognostic importance of glioblastoma features has not been thoroughly explored for sex differences.

**Methods:** Combining multi-modal MR images, biomathematical models, and patient clinical information, this investigation assesses which pretreatment variables have a sex-specific impact on the survival of glioblastoma patients. Pretreatment MR images of 494 glioblastoma patients (299 males and 195 females) were segmented to quantify tumor volumes. Cox proportional hazard (CPH) models and Student’s t-tests were used to assess which variables were associated with survival outcomes.

**Results:** Among males, tumor (T1Gd) radius was a predictor of overall survival (HR=1.027, p=0.044). Among females, higher tumor cell net invasion rate was a significant detriment to overall survival (HR=1.011, p<0.001). Female extreme survivors had significantly smaller tumors (T1Gd) (p=0.010 t-test), but tumor size was not correlated with female overall survival (p=0.955 CPH). Both male and female extreme survivors had significantly lower tumor cell net proliferation rates than other patients (M p=0.004, F p=0.001, t-test). Additionally, extent of resection, tumor laterality, and IDH1 mutation status were also found to have sex-specific effects on overall survival.

**Conclusion:** Despite similar distributions of the MR imaging parameters between males and females, there was a sex-specific difference in how these parameters related to outcomes, which emphasizes the importance of considering sex as a biological factor when determining patient prognosis and treatment approach.

## Introduction

Glioblastoma (GBM) is the most common primary malignant brain tumor, with a median overall survival of 9 to 15 months, depending on the given course of treatment^1-3^, According to Ostrom et al.^4^, only 35% of patients survive more than one year and 4.7% of patients survive more than five years after diagnosis. Factors such as age at diagnosis, Karnofsky performance score (KPS), extent of surgical resection, and tumor location have been found to play a significant role in determining the duration of patient survival^5-7^, but there is still limited insight into which underlying biological features contribute to a patient becoming a “survival outlier.” To date, there is minimal research on the utility of using pretreatment (pre-tx), image-based volumetric and kinetic variables to identify potential extreme and short-term survivors. Additionally, while it has been consistently identified that GBM incidence is higher among males^8-12^ and females GBM patients have better outcomes^8,12-14,^ little to no research has focused on sex-specific predictors of extreme and short-term survival. The ability to pinpoint relevant predictors of the duration of overall survival has clinical value and identifies areas for future research. By using variables derived from patient clinical information and routinely-obtained, non-invasive MR images, we can establish predictors of survival duration that can be readily assessed in a pre-tx setting. Knowing whether these factors affect males and females in the same way will guide research efforts towards best-practice, individualized patient care.

The purpose of this study was to determine whether there are sex-specific predictors of survival outcomes among glioblastoma patients. Using patient data from our multi-institutional brain tumor repository, we tested the significance of eight pre-tx volumetric, kinetic, and clinical variables in predicting extreme and short-term survival. We also tested whether these variables and additional categorical variables, including tumor laterality, extent of resection (EOR), isocitrate dehydrogenase 1 (IDH1) mutation status, and O(6)-methylguanine-DNA methyltransferase promoter (MGMT) methylation status, significantly impacted the overall survival of male and female patients. Throughout the analysis, males and females were tested separately as distinct population groups and their results were compared, allowing us to identify sex-specific impactors of survival outcome among GBM patients.

## Methods

### Imaging

As described in Swanson et al.^15^, tumor volumes were segmented from MR images [gadolinium-enhanced T1-weighted (T1Gd), T2-weighted (T2), and T2 fluid-attenuated inversion recovery (FLAIR)] by trained individuals using our in-house thresholding-based software. These volumes were converted to their spherically-equivalent radii for further analysis.

### Biomathematical Models and Patient-Specific Tumor Kinetics

An extensive literature has been generated over the last two decades applying a biomathematical model to simulate patient-specific glioblastoma growth^15-18^. The primary model is referred to as the Proliferation-Invasion (PI) model and is based on two key parameters: the net rate of proliferation, ϱ, and the net rate of invasion, D. These estimates have been shown to be prognostic of benefit from resection^18^, survival^16^, and radiation efficacy^20^ and can be used to examine therapeutic response^21-22^. Traditional methods of calculating PI D and ϱ require two pre-tx time points of imaging and these are not always available. We have thus leveraged a second model, the Proliferation-Invasion-Hypoxic-Necrotic-Angiogenesis (PIHNA) model^23^, which incorporates necrosis to estimate D and Q using one image time point. For more detail, see **Supplement 16**.

### Patient Population

Our research lab has amassed a large multi-institutional repository consisting of the clinical patient data and serial, multi-modal MR images of over 1400 glioblastoma patients. From this repository, we identified all newly-diagnosed glioblastoma patients with necessary clinical information (sex, age, and overall survival) and a calculated pre-tx (prior to biopsy or resection) tumor volume from a T1Gd MRI. This cohort was comprised of 494 primary GBM patients (299 males and 195 females). Since the calculation of PIHNA D, PIHNA ϱ, and PI D/ϱ requires both T1Gd and T2 or FLAIR (T2/FLAIR) images, a sub-cohort of patients with sufficient imaging was created from the main cohort in order to study the effect of these variables on survival (223 males and 141 females).

We defined extreme survivors (EXS) as those with overall survival (OS) of 5 years (1825 days) or longer. EXS typically make up less than 5% of glioblastoma patients^4^. However, due to the data collection efforts of a multicenter collaboration researching extreme survival among GBM patients (ENDURES), about 9.5% of patients in this cohort were EXS. EXS were compared to Non-EXS (OS<1825 days). We also compared short-term survivors (STS) (OS≤210 days) 24 and Non-STS (OS>210 days). The breakdown of the main cohort and the sub-cohort by sex and survival group is shown in **Table 1**.

**Table 1:** Breakdown of the main cohort and sub-cohort by sex and survival group. Percentages indicate the distribution of males and females in each survival group.

|  | Volumetric and Clinical Data<br>(Main cohort) N=494 |  | PI and PIHNA<br>(Sub cohort)<br>N = 364 |  |
| --- | --- | --- | --- | --- |
|  | Male | Female | Male | Female |
| **All Patients** | 299 (60.5%) | 195 (39.5%) | 223 (61.2%) | 141 (38.7%) |
| **Extreme (OS>1825 days)** | 30 (63.8%) | 17 (36.2%) | 26 (70.3%) | 11 (29.7%) |
| **Short term (OS<210 days)** | 46 (52.3%) | 42 (47.7%) | 32 (50%) | 32 (50%) |

### Statistical analysis

**Table 2** outlines the eight quantitative volumetric, kinetic, and clinical variables that were explored in our investigation. Two-sided Student’s t-tests with Welch’s corrections were used to test whether there were significant differences in the eight quantitative variables between the survival groups. Two-sided Cox-Proportional Hazards models (CPH) were used to assess which of the quantitative variables were significant predictors of OS. Parameters that were significant or almost significant (p<0.10) in univariate analysis were compared in multivariate analysis. Kaplan-Meier survival analysis (two-sided log-rank tests) and CPH models were used to assess the impact of the categorical variables on survival. The following categorical variables were included: IDH1 mutation status, MGMT methylation status, tumor laterality, and EOR. T-tests and Kaplan-Meier survival curves were generated using Prism^25^ and the CPH models were generated using R studio^26^. All statistical analyses were performed separately for the male and female populations. There was no significant difference in the distribution or mean values of these variables between males and females (**Supplement 11**).

**Table 2:** **Definitions and distributions of the eight quantitative volumetric, kinetic, and clinical variables used in this investigation**

| Variable used for Investigation | Definition | Male |  |  | Female |  |  |
| --- | --- | --- | --- | --- | --- | --- | --- |
|  |  | Mean | Median | Range | Mean | Median | Range |
| Age (years) | Age of patient on date of diagnosis | 57.58 | 58 | 12-95 | 58.41 | 60.5 | 9-96 |
| T1Gd Radius (mm) | Combined volume of the central non-enhancing necrotic region and surrounding enhanced region of tumor in a pre-tx T1Gd MR image (converted to a spherically- equivalent radius) | 19.52 | 20.10 | 3.04-33.61 | 19.27 | 18.99 | 4.61-35.08 |
| Necrosis Radius (mm) | Volume of non-enhancing central necrotic region in a pre-tx T1Gd MR image (converted to a spherically- equivalent radius) | 11.39 | 11.69 | 0.00-26.54 | 11.37 | 11.33 | 0.00-27.06 |
| Contrast-enhancing (CE) thickness (mm) | Average linear thickness of the contrast-enhancing region in a pre-tx T1Gd MR image (calculated as the difference between the T1Gd radius and the necrosis radius) | 8.16 | 7.85 | 2.55-18.94 | 7.89 | 7.59 | 0.32-23.26 |
| T2 /FLAIR radius (mm) | Volume of the pre-tx T2 or T2-FLAIR MR image (converted to a spherically- equivalent radius) | 27.11 | 28.31 | 9.94-39.55 | 26.98 | 27.86 | 9.99-42.81 |
| PIHNA D (mm<sup>2</sup>/year) | Net tumor cell diffuse invasion rate | 32.34 | 28.99 | 1.45-145.3 | 36.25 | 23.03 | 0.37-289.9 |
| PIHNA $\rho$ (year<sup>-1</sup>) | Net tumor cell proliferation rate | 65.88 | 18.25 | 1.83-1825 | 82.40 | 18.25 | 1.83-1825 |
| PI D/$\rho$ (mm<sup>2</sup>) | Relative tumor invasiveness | 2.19 | 1.65 | 0.0034-10.26 | 2.12 | 1.28 | 0.0034-10.70 |

### Decision Trees

The decision trees (DT) in this study were created using R^26^, accompanied by a package called *rpar*^28^, which allows effective decision tree pruning. Six DT were produced in total, grouped into 3 pairs. Within each pair, one tree was created using the male population and the other was created using the female population. The PI and PIHNA subcohort of patients (223 males and 141 females) was used to create the training (70% of population) and testing (30%) groups and 10-fold cross validation was used to ensure the generalizability of the results. For each tree, accuracy and sensitivity (EXS and STS are considered condition positive) are reported for the training group, testing group, and the full cohort (training + testing). All six trees were constructed using the eight quantitative pre-tx variables: age, T1Gd radius, necrosis radius, CE thickness, T2/FLAIR radius, PIHNA D, PIHNA ϱ, and PI D/ϱ.

### Study Approval

All featured patients were either consented prospectively or approved for retrospective research before inclusion in this investigation. The usage and collection of patient data was carried out under institutional review board approval.

## Results

### Variables associated with extreme and short-term survival

Student’s t-tests were performed separately on males and females and compared the following groups: EXS vs Non-EXS, EXS vs STS, and STS vs Non-STS. The results of this analysis can be found in Table 3. When compared to the rest of the male population, EXS were significantly younger (p=0.005) and STS were significantly older (p<0.001). Male EXS had significantly smaller Q when compared to male Non-EXS (p=0.004). When compared to the rest of the female population, female EXS were significantly younger (p=0.032) while female STS were significantly older (p<0.001). Female EXS had significantly smaller T1Gd radii compared to female Non-EXS (p=0.010). Compared to the rest of the female population, female EXS had significantly smaller D (p=0.008) and female STS had significantly larger D (p=0.018). Female EXS had significantly smaller ϱ compared to female Non-EXS (p=0.001).

**Table 3:**
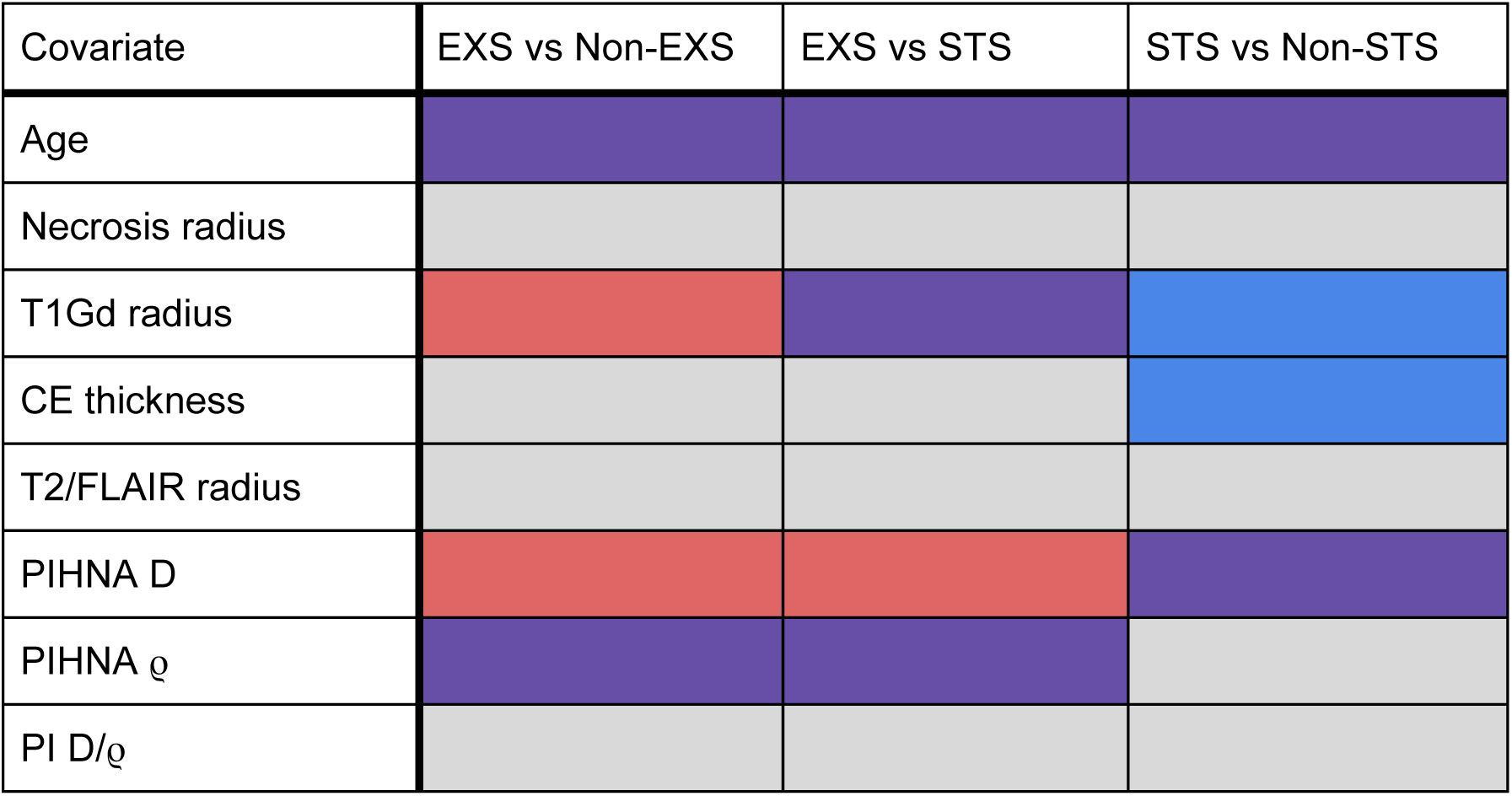
Results of the t-test comparisons of the eight quantitative volumetric and clinical variables between the survival groups for males and females. Purple boxes indicate that the means of the variables were significantly different between the survival groups within both the male and female populations. Red boxes indicate a significant difference within the female population and blue indicate a significant difference within the male population. Gray boxes indicate that neither population showed a significant difference in the means of the variables between the survival groups. Detailed results of t-tests can be found in **Supplement 13**.

In the female EXS vs Non-EXS DT (**Figure 1A and 1B**), the nodes that predicted EXS with 100% sensitivity included T1Gd radius < 21.93 mm and age < 28.5 years. Notably, all male EXS had CE thickness shorter than 11.33 mm, PI D/ϱ above 0.3687 mm^2^, and age below 72 years. In the female EXS vs STS DT (**Figure 1C and 1D**), the nodes that best predicted female EXS included ϱ < 10.33 year ^−1^ and CE thickness < 4.746 mm and the node that best predicted female STS was age ≥ 47.5 years. In the male DT, the node that best predicted EXS was Q < 118.2 year ^−1^ and the node that best predicted STS was D ≥ 11.85 mm^2^/year. The third pair of DT sorted males and females into STS and Non-STS groups (**Figure 1E and 1F**). Among females, the nodes that best predicted STS included age ≥ 49.5 years, T2/FLAIR radius ≥ 23.76 mm, and D ≥ 41.23 mm^2^/year. In the male DT, the nodes that most accurately predicted STS included age ≥ 47.5 years, ϱ ≥ 10.33 year ^−1^, and CE thickness between 11.25 mm and 12.36 mm.

**Figure 1:**
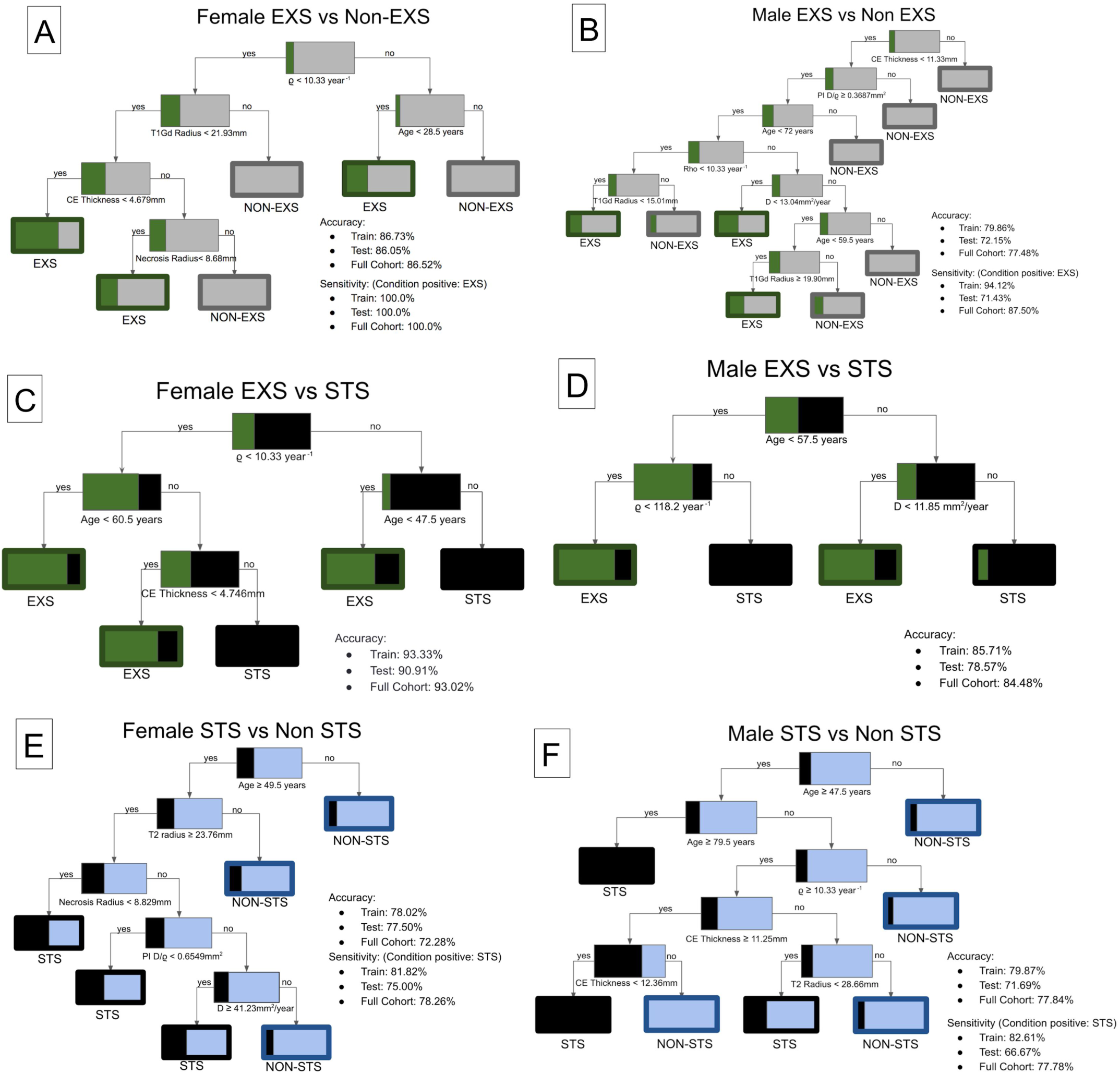
Decision trees separating male and female EXS, Non-EXS, STS, and Non-STS. At each node, color (green for EXS, gray for Non-EXS, black for STS, and blue for Non-STS) and percentages indicate concentration of each group. Percentages in red indicate concentration of IDH1 mutant patients at each endpoint. **A)** Female EXS vs Non-EXS DT (n=141). **B)** Male EXS vs Non-EXS DT (n=223). **C)** Female EXS vs STS DT (n=43). **D)** Male EXS vs STS (n=58). **E)** Female STS vs Non-STS DT (n=141). **F)** Male STS vs Non-STS DT (n=223).

### Variables associated with overall survival

Univariate and multivariate CPH analyses **(Table 4)** were utilized to determine which variables significantly influenced the overall survival of GBM patients. Variables that were significant or almost significant (p<0.10) in univariate analysis were analyzed in multivariate analysis. In the male multivariate CPH, factors found to independently influence survival included: age (HR=1.030, p<0.001) and T1Gd radius (HR=1.027, p=0.044). In the female multivariate CPH analysis, age (HR=1.021, p=0.006) and PIHNA D (HR=1.011, p<0.001) were identified as significant independent prognostic factors.

**Table 4:** Results of univariate and multivariate CPH analyses for males and females. Factors that were almost significant (p<0.10) or significant in univariate analysis were included in the multivariate analysis.

| **Males** | **Univariate** |  |  | **Multivariate** |  |  |
| --- | --- | --- | --- | --- | --- | --- |
| Covariate | HR | 95% CI | p-value | HR | 95% CI | p-value |
| Age | 1.027 | 1.018-1.037 | **<0.001** | 1.030 | 1.017-1.044 | **<0.001** |
| Necrosis radius | 1.018 | 0.996-1.040 | 0.118 |  |  | N/A |
| T1Gd radius | 1.024 | 1.003-1.046 | **0.025** | 1.027 | 1.001-1.054 | **0.044** |
| CE Thickness | 1.028 | 0.989-1.068 | 0.161 |  |  | N/A |
| T2/FLAIR radius | 0.996 | 0.972-1.020 | 0.744 |  |  | N/A |
| PIHNA D | 1.003 | 0.997-1.010 | 0.266 |  |  | N/A |
| PIHNA $\rho$ | 1.001 | 1.000-1.001 | 0.064 | 1.000 | 0.999-1.001 | 0.637 |
| PI D/$\rho$ | 0.932 | 0.872-0.996 | **0.038** | 0.951 | 0.880-1.029 | 0.210 |

| **Females** | **Univariate** |  |  | **Multivariate** |  |  |
| --- | --- | --- | --- | --- | --- | --- |
| Covariate | HR | 95% CI | p-value | HR | 95% CI | p-value |
| Age | 1.028 | 1.015-1.041 | **<0.001** | 1.021 | 1.006-1.037 | **0.006** |
| Necrosis radius | 1.017 | 0.991-1.042 | 0.204 |  |  | N/A |
| T1Gd radius | 1.026 | 1.000-1.052 | **0.048** | 0.993 | 0.964-1.023 | 0.641 |
| CE Thickness | 1.037 | 0.988-1.088 | 0.143 |  |  | N/A |
| T2/FLAIR radius | 1.017 | 0.989-1.045 | 0.232 |  |  | N/A |
| PIHNA D | 1.011 | 1.006-1.016 | **<0.001** | 1.011 | 1.005-1.017 | **<0.001** |
| PIHNA $\rho$ | 1.001 | 1.000-1.002 | 0.052 | 1.000 | 0.999-1.002 | 0.801 |
| PI D/$\rho$ | 0.996 | 0.937-1.059 | 0.906 | | | N/A |

### IDH1 Mutation

Since IDH1 mutation has been previously identified as significant predictor of long-term survival^14^, we analyzed the impact of sex and IDH1 status on the overall survival of our patient cohort. 120 patients in the main cohort had determined IDH1 status, consisting of 69 wild-type (wt) and 8 mutant (mut) male patients and 39 wt and 4 mut female patients. When looking at the entire population (both males and females), there was a trend towards IDH1 mut patients having better survival (log-rank, p=0.071). Among females, IDH1 mut survived significantly longer than IDH1 wt patients (log-rank, p=0.008), but among males, the survival difference was not significant (log-rank, p=0.924) **(Supplement 1)**. All 4 IDH1 mut females survived at least three years, making them all long-term survivors^29^.

We also assessed whether IDH1 mut patients had the same features as the extreme survivors in this analysis (younger age, lower PIHNA D, lower PIHNA ϱ, and smaller T1Gd radii). Unlike the female EXS, IDH1 mut females did not have lower PIHNA D (t-test, p=0.402) or smaller T1Gd radii (p=0.584) compared to their wt counterparts, but they did have significantly lower PIHNA ϱ when compared to wt females (p=0.027). Males did not show significantly different PIHNA D (p=0.796) or PIHNA ϱ (p=0.461) between the two IDH1 status groups, but IDH1 mut males did tend to have smaller T1Gd radii (p=0.052) when compared IDH1 wt males. Both male and female IDH1 mut were significantly younger than their wt counterparts (Male p=0.024, Female p=0.007).

### MGMT Methylation

Methylation of the MGMT promoter has been found to be more common in long-term survivors^30^, so we also assessed the impact of MGMT methylation on the survival of our population cohort. Ninety patients from the main cohort had available MGMT methylation status, which comprised of 32 females (12 methylated and 20 unmethylated) and 58 males (18 methylated and 40 unmethylated). Methylated patients had significantly better survival than unmethylated patients among males (log-rank, p=0.013), females (p=0.007), and the entire population (males and females) (p<0.001) **(Supplement 4)**. Multivariate CPH analyses that assessed the impact of MGMT status on survival while accounting for age showed that MGMT status significantly impacted survival for males (p=0.004) and females (p=0.037). Among EXS with available MGMT methylation status (n=15), 50% (n=5) of males and 60% (n=3) of females had MGMT methylation, while among Non-EXS (n=75), 29% (n=14) of males and 33% (n=9) of females had MGMT methylation, suggesting that MGMT methylation was more common among both male and female EXS.

When we tested to see if MGMT methylated patients shared the features of extreme survivors (younger age, lower PIHNA D, lower PIHNA ϱ, and smaller T1Gd radii), we found that MGMT methylated females had significantly lower ϱ (t-test, p=0.026) and tended to have lower D (p=0.057) when compared to MGMT unmethylated females. There was no significant difference in the values of D (p=0.477) or ϱ (p=0.869) between MGMT methylated and unmethylated males. For both males and females, there was no significant difference in age (Male p=0.724, Female p=0.735) or T1Gd radii (Male p=0.397, Female p=0.241) between methylated and unmethylated patients.

### Laterality

Using pre-tx T1Gd MR images, we determined the laterality of each patient’s tumor, classifying the tumors as being located in the right hemisphere, left hemisphere, or both hemispheres (bilateral). The impact of tumor laterality on survival was assessed separately for males and females, and the results were compared. Among males, there were 129 left hemisphere GBMs, 154 right hemisphere GBMs, and 11 bilateral GBMs, and among females there were 86 left hemisphere GBMs, 96 right hemisphere GBMs, and 9 bilateral GBMs. Laterality could not be determined for 5 male and 4 female patients.

Male patients with tumors on the left side tended to have better survival than males with tumors on the right side (log-rank, p=0.077) and had significantly better survival than males with bilateral tumors (p=0.010) **(Supplement 6)**. In a multivariate CPH analysis that also accounted for extent of resection, tumor location in the left hemisphere was found to be a significant independent predictor of improved survival outcome for males (p=0.017) **(Supplement 14)**. There were more EXS than STS among males with tumors on the left side and there were almost twice as many STS as EXS among males with tumors on the right side. Laterality did not have a significant impact on survival for female patients (CPH, p=0.299) **(Supplement 14)**. There was no significant difference in survival between females with left and right hemisphere tumors (log-rank, p=0.218), and females with bilaterally located tumors did not have significantly worse survival when compared to females with non-bilateral tumors (bilateral vs left p=0.272, bilateral vs right p=0.471) **(Supplement 6)**.

### Extent of Resection

Our investigation evaluated whether the extent of initial surgical intervention, a known prognostic factor among GBM patients, had the same prognostic value for both male and female GBM patients. Patient EOR status, categorized as gross total resection (GTR), subtotal resection (STR), or biopsy, was obtained from the patient records. From the main cohort of 494 patients, 211 males (83 GTR, 83 STR, and 45 biopsy) and 136 females (54 GTR, 55 STR, and 27 biopsy) had available EOR status.

EOR had a significant impact on the survival of male GBM patients. GTR males had significantly better survival than STR males (log-rank, p=0.033) **(Supplement 9)** and males who received some surgical resection (GTR or STR) had significantly better survival than males who only received a biopsy (p=0.013) **(Supplement 8)**. Cochran-Armitage Trend Test showed that there was significant trend towards male EXS receiving more extensive resections and male STS receiving less extensive resections or biopsies (p=0.027). Female who received resection (GTR or STR) trended towards improved survival compared to biopsy females (log-rank, p=0.077) **(Supplement 8)**, but there was no significant difference in survival between GTR females and STR females (p=0.992) **(Supplement 9)**. Additionally, EOR did not significantly impact female survival in univariate CPH analysis (p=0.180) **(Supplement 14)**. Trend test showed that there was an insignificant trend towards female EXS receiving more extensive resections and female STS receiving less extensive resections or biopsies (p=0.098).

### Patients receiving current standard of care

Due to the timespan over which they were collected, the patients in our cohort received a wide variety of treatment protocols. In order to ensure that our results maintain significance among patients who receive the current standard of care (maximal safe resection followed by concurrent temozolomide and radiation therapy), we created a subset of patients who received this treatment protocol (Stupp protocol patients)^32^ and tested which factors were associated with overall survival among those patients **(Supplement 15)**. In this limited subpopulation, we had 113 males and 66 females **(Supplement 15A)**. Among females, PIHNA D was a significant independent predictor of overall survival and among males, PIHNA ϱ was a significant independent predictor of overall survival **(Supplement 15B)**.

## Discussion

While there are no differences in the distributions of these quantitative and categorical variables between males and females, this investigation found that there are sex-specific differences in the impact that these variables have on patient survival **(Figure 2)**.

**Figure 2:**
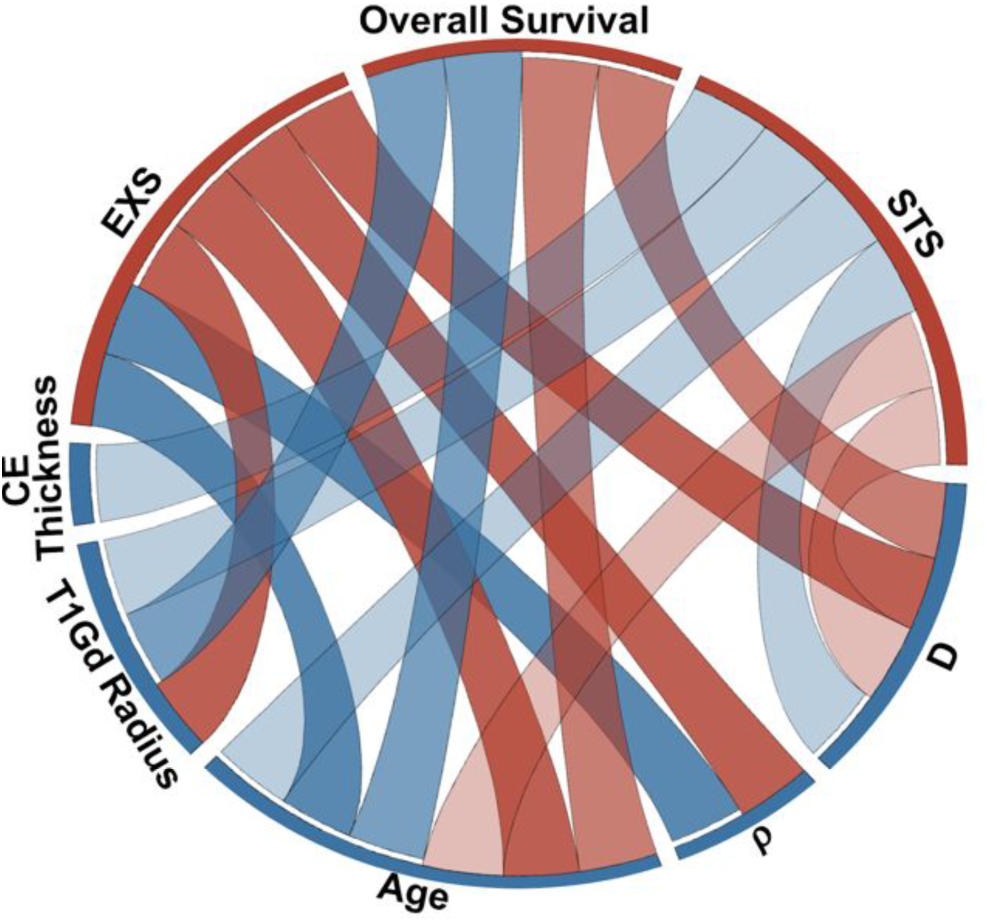
Sex differences in association with EXS, STS, or overall survival^31^. The bottom portion of the outer ring shows the relevant quantitative variables and the top portion shows the three aspects of survival that are associated with these variables (EXS, STS, and Overall Survival). Red ribbons indicate significant relationships for female patients and blue ribbons indicate significant relationships for male patients. Variables that were significant in multivariate CPH are connected to the Overall Survival segment and variables that were significant in Student t-tests with Welch’s correction are connected to the relevant EXS or STS segments.

### Impact of quantitative variables on survival

Among females, tumor cell diffuse invasion rate (PIHNA D) is strongly negatively correlated with overall survival for females across the various analyses. Notably, both when EOR was included in multivariate CPH analysis **(Supplement 14)** and when only Stupp protocol patients were considered **(Supplement 15B)**, PIHNA D was still an independent predictor of survival for females. Although it was not significant in the CPH multivariate analysis, it is notable that males had a significant positive association between overall survival and PI D/ϱ in univariate analysis **(Table 4)**. This suggests that more nodular tumors at time of diagnosis are associated with worse prognosis for males, which is contrary to the finding that more diffusely invasive tumors are associated with worse prognosis for females.

Smaller total tumor size (T1Gd radius) is significantly associated with EXS for females. DT analysis showed that nodes isolating females with below average necrosis radii and CE thickness, both components of overall tumor size, were highly sensitive predictors of EXS **(Figures 1A and 1C)**.When the mean T1Gd radius of EXS was compared to the mean T1Gd radius of other survival groups, the mean radius of EXS was significantly smaller **(Table 3)**. Univariate CPH found that T1Gd radius size was a significant predictor of survival **(Table 4)**, but if EXS were excluded from the analysis, this relationship is no longer significant (p=0.503). These results suggest female extreme survivors have smaller pre-tx T1Gd radii, but T1Gd radius is not negatively correlated with overall survival for females in general.

Among males, total tumor size (T1Gd radius) is negatively correlated with overall survival across the statistical analyses **(Tables 3 and 4)**. In the DT analyses, CE thickness, a component of total tumor size, is a highly sensitive predictor of survival outcome **(Figures 1B and 1F)**. It is notable total tumor size at time of diagnosis is negatively correlated with outcome for males and not females.

Age is known to have a significant impact on the survival of glioblastoma patients^5-7^ and this analysis confirmed that age significantly impacts the survival of both males and females. Across the analyses, older age at time of diagnosis is consistently associated with shorter survival, while younger age is associated with longer survival **(Table 3 and 4)**.

Lower tumor cell proliferation rates (PIHNA ϱ) are associated with EXS for both males and females. DT analysis and statistical analysis both showed that low proliferation rates were associated with EXS **(Table 3 and Figures 1C and 1D).** Low tumor cell proliferation rates appear to be predictive of long-term survival for both males and females, but high rates do not appear to predict short-term survival.

### Impact of categorical variables on survival

While Schiffgens et al.^34^ found that only IDH1 mutant males demonstrate significantly improved survival compared to IDH1 wild-type males, our investigation found the opposite, that only IDH1 mutant females demonstrate significantly improved survival when compared to their wild-type counterparts **(Supplement 1)**. While our study does have a relatively small sample of IDH1 mutants, our finding is in concurrence with the findings of Yang et al.^33^, who grouped females by genetic similarities and found that the longest-living female cohort predominantly consisted of IDH1 mutant females. They did not see this effect for males. Our IDH1 mutant females were all long-term survivors and they demonstrated the same depression in PIHNA ϱ when compared to the wild-type females that the EXS females demonstrated when compared to Non-EXS females. However, IDH1 mutant females did not have lower PIHNA D compared to the wild-type population. Meanwhile, IDH1 mutant males did not show improved survival, depressed PIHNA ϱ, or significantly different PIHNA D when compared to IDH1 wild-type males. In Baldock et al.^17^, IDH1 mutation was shown to be significantly correlated with lower ϱ and higher D/ϱ (lower ϱ/D) among contrast-enhancing glioma patients. The sexes were not separated in this analysis, so there is a possibility that the effect of the depressed ϱ may have only existed for females. The findings of Schiffgens et al.^34^ Yang et al.^33^, and this investigation make a compelling case for the need to consider sex in IDH1-related research.

Previous studies have demonstrated that MGMT promoter methylation is a significant independent prognostic factor^37^ and is more common among long-term survivors^30,38^. Despite having a relatively small sample of patients with known MGMT methylation status, our analysis was able to confirm that, for both males and females, MGMT methylation was more common among extreme survivors and was a significant independent prognostic factor. Previous studies have also found that the survival benefit of MGMT methylation was stronger or only significant among female patients^34,39^, but our analysis did not see any evidence of females benefiting more from MGMT methylation than males. However, our analysis did show that methylated females had some of the same characteristics as extreme surviving females, namely that methylated females had lower PIHNA D and significantly lower PIHNA ϱ when compared to unmethylated females.

In this investigation, GBM laterality impacted male survival, but had no impact on female survival. Even after accounting for EOR, males with tumors located in the left hemisphere had a significant survival advantage compared to males with tumors located in the right hemisphere. Ellingson et al.^40^ found that patients who responded favorably to chemotherapy, patients with prolonged survival, and patients with specific genetic modifications, like MGMT promoter methylation and IDH1 mutation, had tumors that clustered in areas of the left hemisphere of the brain. Additional research will need to be conducted on the relationship between genetic modifiers, laterality, sex, and survival.

Previous literature has identified extent of resection as a significant predictor of overall survival for GBM patients^6,18,41-42^, but whether EOR has the same impact on survival for males and females has not been clearly elucidated. Our analysis found that EOR has a significant impact on the survival of male GBM patients, with a more complete resection being associated with longer survival and potentially extreme survival. Among females, there was a survival benefit associated with receiving resection, but the extent of resection did not have a significant impact on survival. These findings suggest that EOR may have a sex-specific impact on survival, but further study will be required to fully understand the extent of this difference.

### Limitations and Further Work

Due to the utilization of retrospective clinical data, it was not possible to control for all confounding factors and bias within our dataset. However, our utilization of a large cohort of almost 500 patients allows for the mitigation of some of these confounding effects. The findings presented in this investigation lay the groundwork for future research on the topic of sex differences in prognostic indicators of extreme survival in patients with GBM. Shinojima et al.^8^ observed that their cohort of extreme survivors consisted entirely of females and had a disproportionately large number of giant cell glioblastoma cases. Future work could consider whether histological variations in GBM have sex-specific effects on survival. Additionally, considering more sensitive and individualized elements of the tumor, like the biological environment surrounding the tumor, could provide a more thorough understanding of what makes survival outliers unique.

## Conclusion

Taken together, these results further validate the need to consider sex as a relevant biological factor in all glioblastoma-related research. Sex has been shown to significantly impact GBM incidence and prevalence^8-11^, survival^8,13-14^, oncogenic gene expression^33^, glycolytic pathway gene expression^43^, and now the predictors of overall survival. Despite these findings, many studies do not specify patient sex and those that do often do not consider sex when reporting the results of their analysis. The consideration of the role of sex in tumor behavior, incidence, growth, and treatment response will only lead to higher-quality, more individualized knowledge and care for glioblastoma patients.

